# TRIM69 inhibits Vesicular Stomatitis Indiana Virus (VSIV)

**DOI:** 10.1101/669176

**Authors:** Suzannah J. Rihn, Muhamad Afiq Aziz, Douglas G. Stewart, Joseph Hughes, Matthew L. Turnbull, Mariana Varela, Elena Sugrue, Christie S. Herd, Megan Stanifer, Steven P. Sinkins, Massimo Palmarini, Sam J. Wilson

**Affiliations:** MRC-University of Glasgow Centre for Virus Research, Glasgow, United Kingdom; Department of Infectious Diseases, Virology, Heidelberg University Hospital, Heidelberg, Germany

## Abstract

Vesicular Stomatitis Indiana Virus (VSIV) is a model virus that is exceptionally sensitive to the inhibitory action of interferons. Interferons induce an antiviral state by stimulating the expression of hundreds of interferon stimulated genes (ISGs). These ISGs constrain viral replication, limit tissue tropism, reduce pathogenicity and inhibit viral transmission. Because VSIV is used as a backbone for multiple oncolytic and vaccine strategies, understanding how ISGs restrict VSIV, not only helps in understanding VSIV-pathogenesis, but helps evaluate and understand the safety and efficacy of VSIV-based therapies. Thus there is a need to identify and characterize the ISGs that possess anti-VSIV activity. Using arrayed ISG expression screening, we identified TRIM69 as an ISG that potently inhibits VSIV. This inhibition was highly specific as multiple viruses (including influenza A virus, HIV-1, Rift Valley Fever Virus and dengue virus) were not affected by TRIM69. Indeed, just one amino acid substitution in VSIV can govern sensitivity/resistance to TRIM69. TRIM69 is highly divergent in human populations and exhibits signatures of positive selection that are consistent with this gene playing a key role in antiviral immunity. We propose that TRIM69 is an IFN-induced inhibitor of VSIV and speculate that TRIM69 could be important in limiting VSIV pathogenesis and might influence the specificity and/or efficacy of vesiculovirus-based therapies.

**IMPORTANCE:** Vesicular Stomatitis Indiana Virus (VSIV) is a veterinary pathogen that is also used as a backbone for many oncolytic and vaccine strategies. In natural and therapeutic settings, VSIV infection is sensed by the host and host-cells make proteins that protect them from viruses. In the case of VSIV, these antiviral proteins constrain viral replication and protect most healthy tissues from virus infection. In order to understand how VSIV causes disease and how healthy tissues are protected from VSIV-based therapies, it is crucial that we identify the proteins that inhibit VSIV. Here we show that TRIM69 is an antiviral defence that can potently and specifically block VSIV infection.

## INTRODUCTION

Most invading pathogens are sensed by the vertebrate host ensuring that immune defences are deployed appropriately. Following sensing, one common outcome is the secretion of type I interferons (IFNs) whose signalling results in the upregulation of hundreds of IFN stimulated genes (ISGs) (1, 2). Many ISG products interfere with viruses directly, generating an ‘antiviral state’ in stimulated cells that impedes the infection, replication or propagation of viruses (3–5). In addition, many ISGs are themselves involved in pathogen sensing and signal transduction, placing cells in a heightened state of alert poised to detect invading pathogens (6). The IFN response typically involves hundreds of ISGs, many of whom have been regulated by IFNs for hundreds of millions of years (1). Although the major role that IFNs play in constraining viral pathogenesis and viral colonization is well-established, because IFN responses involve so many ISGs, it is often unclear which individual gene-products inhibit a given virus.

Vesicular stomatitis Indiana virus (VSIV) is a virus that is mainly restricted to the Americas where it causes vesicular stomatitis, a disease that primarily affects ungulates and rarely causes mild infections in humans (7–9). VSIV is transmitted by biting insects and causes characteristic vesicular lesions at bite sites around the hooves, mouth, nose, teats and coronary bands (7). Although complications can occur, natural VSIV infection is typically mild and rapidly resolved. In contrast, experimental VSIV infection can be highly pathogenic and neurotropic in young mice (10).

In addition to being a notable veterinary pathogen, VSIV has been used extensively as a model virus and has been integral to our understanding of vesiculovirus and rhabdovirus biology. Notably, vesiculoviruses can be particularly sensitive to IFNs, leading to their inclusion in the IFN unit definition assay (11). Indeed, type I IFNs likely play a major role in limiting the severity of VSIV infections and multiple ISGs have been ascribed anti-VSV activity (4, 12–15).

Importantly, IFNs play a major role in constraining VSIV *in vivo*. Type I IFN receptor (IFNAR) KO mice succumb to doses of VSIV that are several orders of magnitude lower than a lethal dose in wild-type (WT) mice (16). Moreover, whilst VSIV is largely restricted to the central nervous system (CNS) in lethally infected WT mice, VSIV colonizes multiple organs in IFNAR KO mice (16). Interestingly, it is likely that multiple ISGs are involved in limiting the tissue tropism of VSIV. Specifically, it appears that IFIT2 is crucial for preventing VSIV colonization of the brain but it is not solely responsible for limiting VSIV replication in other organs (15). Thus, other ISGs must play key roles in limiting VSIV tissue tropism. Importantly, VSIV causes neurological disease in multiple species following intracranial inoculation (17, 18), suggesting that the ability of ISGs to prevent VSIV from initially accessing the CNS is the cornerstone in limiting VSIV neuropathology across multiple species (19).

VSIV’s low pathogenicity in humans, its rapid replication, and ease of genetic manipulation have made this virus the basis of multiple therapeutic strategies. For example, VSIV can be modified to express antigens from heterologous viruses that can be utilised as vaccine strategies (20). This approach has achieved recent notable success in conferring protection from Ebola virus infection (21). Similarly, VSIV has been used as the backbone of multiple oncolytic strategies (22). Just like natural VSIV infection, IFNs and ISGs appear to be critical for preventing oncolytic viruses from invading healthy tissues (23, 24) and could be critical determinants governing whether oncolytic vesiculoviruses will be efficacious (25). Furthermore, ISGs likely play a key role in limiting the replication of VSIV-based vaccines and are an important safety feature of this immunization strategy.

The key roles that ISGs play in constraining VSIV pathogenesis and limiting VSIV replication (in natural infection, oncolytic therapies and vaccine strategies) means there is a need to better understand how ISGs inhibit VSIV. Using arrayed ISG expression screening, we identified that TRIM69, a relatively poorly characterised TRIM protein, has anti-VSIV activity. Through exogenous expression and CRISPR Cas9 knockout, we demonstrate that both exogenous and endogenous TRIM69 have potent anti-VSIV activity. Importantly, the inhibition is highly specific for VSIV and multiple other viruses were not inhibited by TRIM69. Notably, TRIM69 shows strong signatures of positive selection and multiple common alleles circulate in human populations. Interestingly, murine orthologues of TRIM69 had no detectable anti-VSIV activity whereas rat TRIM69 possessed potent antiviral activity. We speculate that TRIM69 could be an important ISG for protecting healthy tissues from VSIV and might therefore limit VSIV pathogenesis and influence the specificity and efficacy of vesiculovirus-based therapeutic strategies.

## RESULTS

### ISG expression screening reveals the anti-VSIV activity of TRIM69

We have previously used arrayed ISG expression screening of human and rhesus macaque ISG libraries to identify antiviral factors targeting a range of viruses (3, 5, 26). Although, VSIV has previously been subjected to a large-scale screen of ~300 interferon induced genes (4), we reasoned that using larger libraries of arrayed ISGs might identify additional anti-VSIV effectors. We recently expanded our human ISG library to include >500 ISGs, which can be used in conjunction with our existing library of >300 rhesus macaque ISGs (5, 26) (Figure 1A), all of which are encoded by lentiviral vectors (Figure 1B). In addition, we took advantage of a single-cycle VSIV-GFP system (rVSVΔG-GFP, referred to herein as VSIV-GFP) to allow us to identify strong early blocks to VSIV with high fidelity (27). We first transduced human MT4 cells with each ISG-encoding lentiviral vector and then challenged these cells with VSIV-GFP (using a dose where ~30% of cells were infected). The level of VSIV-GFP infection in the presence of each individual ISG was then quantified using flow cytometry (Figure 1D). Strikingly, only three genes potently inhibited VSIV under these conditions: macaque IFNB1, human Mx1 and human TRIM69. Although we identified only one ortholog of each gene, we do not ascribe this to species-specific antiviral activity as macaque TRIM69 and human IFNB1 were not present in these libraries. Moreover, the isoform of macaque Mx1 included in the screen lacked 154 N-terminal residues relative to the human counterpart (that exhibited anti-VSIV activity), potentially explaining the lack of inhibition conferred by the macaque variant. Importantly, TRIM69 was not identified in the previous ISG screen of VSIV, as it was not present in the ISG library used (4).

**Figure 1.**
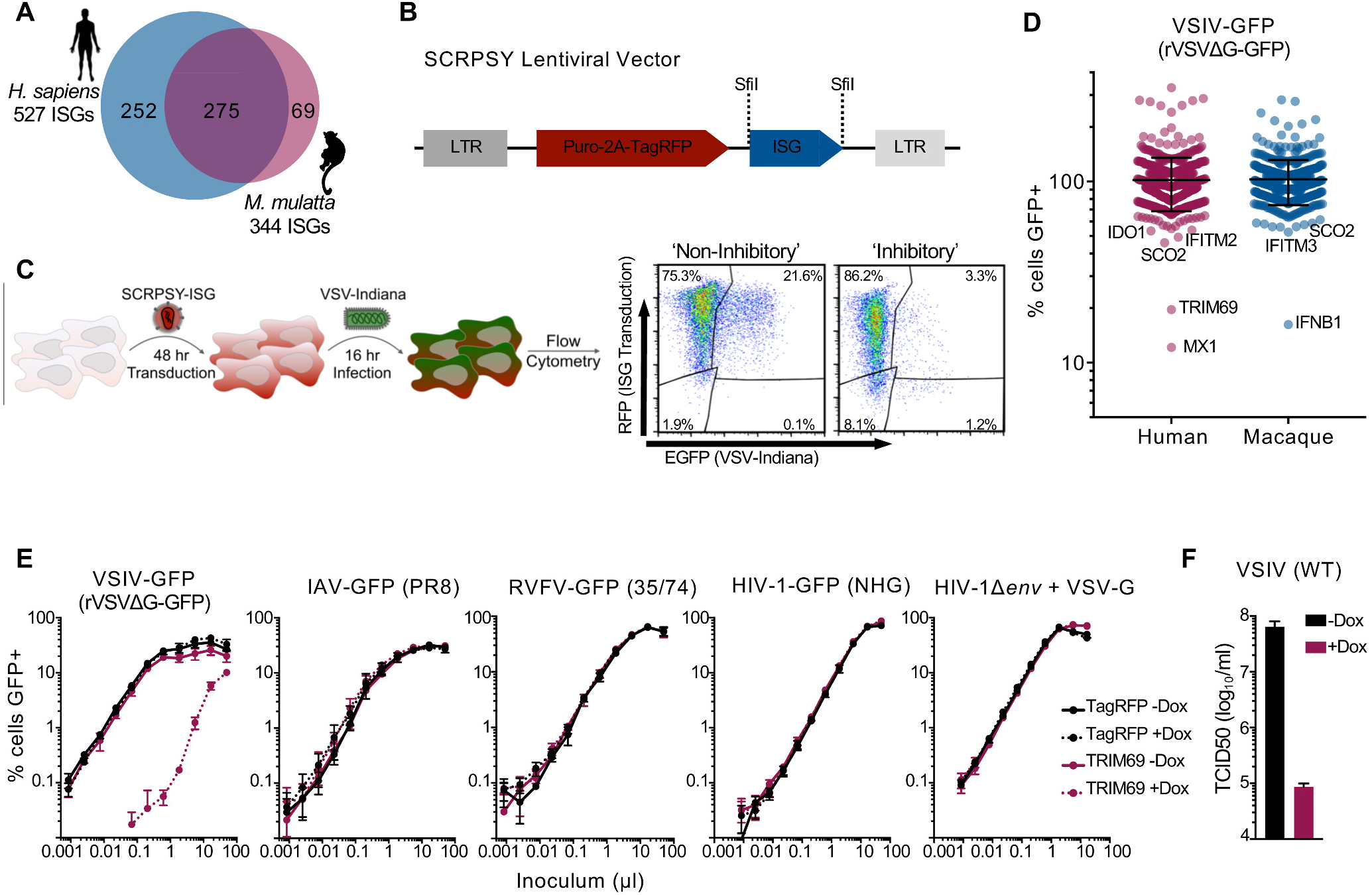
Arrayed ISG expression screening reveals the potent anti-VSIV activity of TRIM69. (A) a schematic of the lSG libraries used herein. (B) a schematic of the SCRPSY lentiviral vector (KT368137.1) used to deliver ISGs in C and D (one ISG per well of a 96-well plate). (C) A schematic of the ISG screening pipeline used in D. (D) Normalised infection (median centred) of cells expressing different ISGs (each dot represents the observed infection in the presence of a single ISG). The screen was executed once. (E) MT4 cells (modified to express doxycycline inducible TRIM69) were incubated with and without doxycycline for 24 h and then challenged with serially diluted GFP-encoding variants of VSIV for 16 h, IAV (influenza A virus) for 16 h, RVFV (Rift Valley Fever phlebovirus) for 48 h and HIV-1 (Human Immunodeficiency Virus) for 48 h prior to fixation and enumeration of GFP positive cells using flow cytometry. virus titrations were carried out on at least 2 occasions and typical results are shown. Mean values of experimental replicates are plotted and error bars represent standard deviation. (F) The same cells as in E were infected with unmodified (WT) VSIV (and infectivity/replication was quantified using TCID50) The mean and standard deviation are plotted.

### TRIM69 exhibits potent and highly specific antiviral activity

Because the anti-VSIV activity of type I IFN and Mx1 are well documented (13, 16), we were immediately struck by how potent the TRIM69-mediated inhibition of VSIV was in our initial screen (Figure 1D). At the time that these experiments were carried out, TRIM69 had not been ascribed any antiviral activity, so we examined the ability of doxycycline inducible TRIM69 to inhibit a small panel of viruses. Notably, while VSIV-GFP infection was reduced by >100-fold by TRIM69, the other viruses in our panel (Influenza A (PR8 (A/Puerto Rico/8/1934 (H1N1)), RVFV (35/74) and HIV-1 (NHG)) were unaffected by TRIM69 expression (Figure 1E). It is well documented that many TRIM proteins are involved in antiviral signalling, which can often be triggered by exogenous expression (5, 28–30). Moreover, exogenous or endogenous expression of many ISGs can promote cell death (31). However, the highly specific antiviral activity of TRIM69, suggests that the antiviral mechanism does not involve global processes such as cellular toxicity or the induction of a polygenic antiviral state. Interestingly, a VSV-G (glycoprotein) pseudotyped variant of HIV-1, that does not express an HIV-1 envelope glycoprotein, was also insensitive to TRIM69-inhibition. This suggests that VSIV-inhibition occurs after viral entry and that the VSIV glycoprotein is not directly targeted by TRIM69. Importantly, unmodified wild-type replication competent VSIV, was also potently inhibited by TRIM69 (Figure 1F).

### Endogenous TRIM69 is IFN-inducible and potently inhibits VSIV

We have previously used comparative transcriptomics to study the IFN response in a variety of species (1). Meta-analysis of these data indicated that TRIM69 expression was upregulated ~10-fold, following IFN-stimulation of primary human fibroblasts (Figure 2A). Similarly, TRIM69 has previously been identified as an ISG in multiple studies, and is typically induced between ~2 and ~10-fold by type I IFNs (2, 32). To examine whether the endogenous protein exhibited antiviral activity, we knocked out TRIM69 using CRISPR/Cas9 in diploid CADOES1 cells (33). TRIM69 knock out (KO) single-cell clones were derived and the KO was confirmed by sequencing the genetic lesions. In accordance with the IFN-sensitive nature of vesiculoviruses, IFN treatment potently blocked VSIV infection in “no guide” control clones (by ~25000 fold) (Figure 2B). In striking contrast, when TRIM69 was knocked out, the protective effect of IFN was markedly reduced (by ~300 fold). We interpret this as endogenous TRIM69 playing a major role in the anti-VSIV effects of IFN. Once again, the TRIM69 anti-VSIV activity was specific, as the magnitude of HIV-1 inhibition was similar in the presence or absence of TRIM69 (Figure 2C).

**Figure 2.**
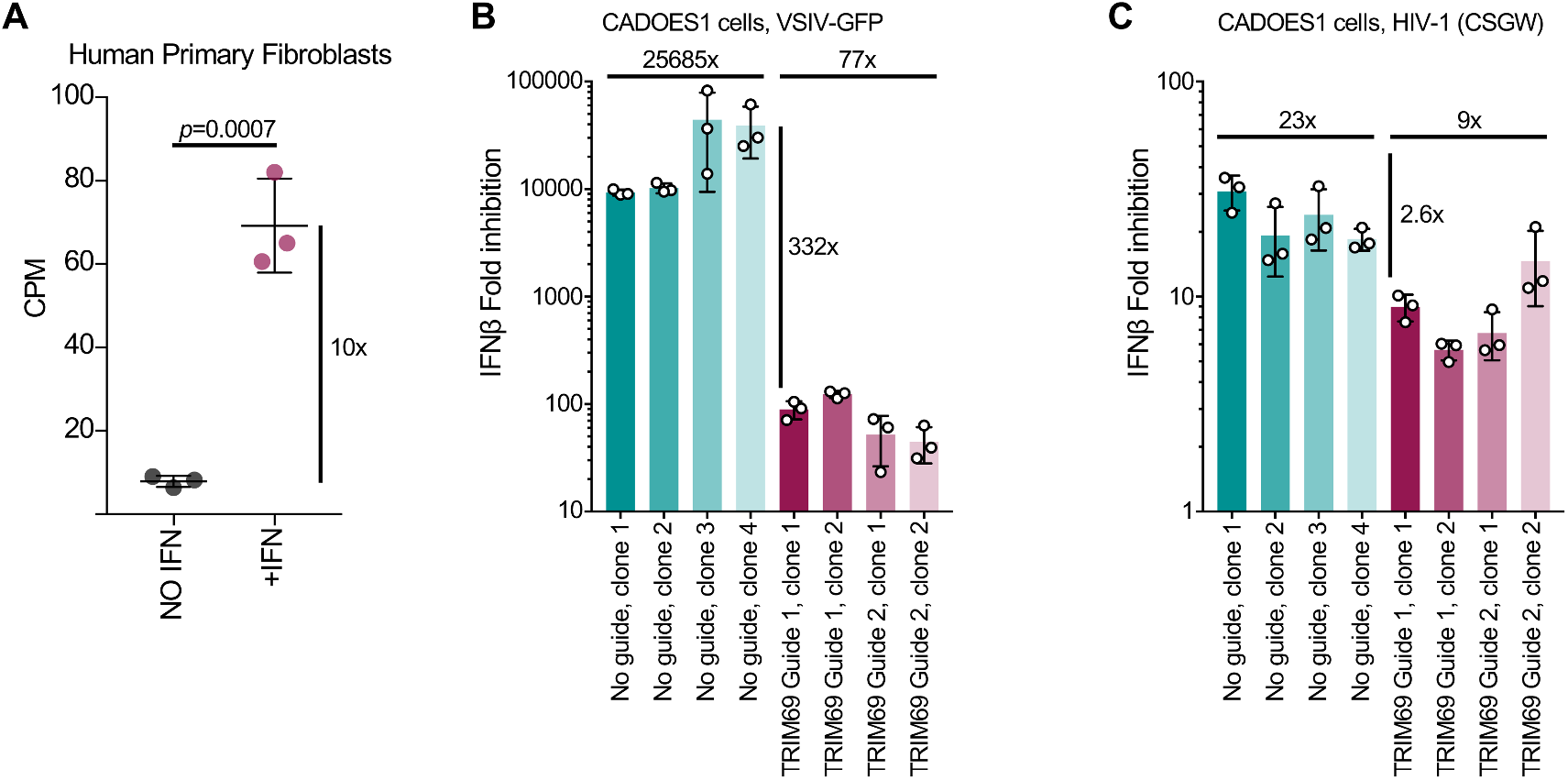
IFN stimulated Endogenous TRIM69 has potent anti-VSIV activity. (A) TRIM69 expression in primary human fibroblasts following 4 h in the presence or absence of 1000 units/ml of universal IFN. The transcriptomes were defined using RNAseq (1) and the meta-analyzed TRIM69 expression is plotted. (B) VSIV-GFP inhibition was measured in CADOES1 cells where TRIM69 expression was knocked out using CRISPR CAS9 or in transduced ‘no guide’ controls. Titrated challenges were used to determine the titre and calculate the fold-inhibition16 h after infection. Fold-inhibition was calculated from 3 independent experiments, each experiment is represented by the circles and the mean and standard deviation is plotted. (C) As in B using a single cycle HIV-1 reporter system (CSGW) (65).

#### Not all TRIM69 isoforms confer antiviral activity

Because alternative splicing can produce divergent variants of antiviral factors, these spliced isoforms can exhibit differential antiviral activity. In the case of TRIM5, the spliced isoforms have been informative in understanding the mechanism of TRIM5’s antiviral activity (34). We therefore considered whether all isoforms of TRIM69 conferred anti-VSIV activity. We cloned the five TRIM69 isoforms listed on ENSEMBL and we considered their activity in our doxycycline inducible system. Under these conditions, the only variant that conferred anti-VSIV activity was the longest isoform, isoform A (Figure 3A-F), which potently inhibited VSIV-GFP infection. One caveat is that (despite multiple attempts) we were unable to verify the expression of isoforms C, D and E (Figure 3G). We cannot therefore distinguish between technical difficulties (such as the antibody not recognising these isoforms) or biological processes (such as the rapid turnover of these isoforms) in preventing us from visualizing these proteins. Nevertheless, we were able to detect abundant expression of isoform B (Figure 3G) and conclude that this isoform has no anti-VSIV activity. This suggests that the RING domain is critically required for anti-VSIV activity or the correct folding or multimerization of TRIM69.

**Figure 3.**
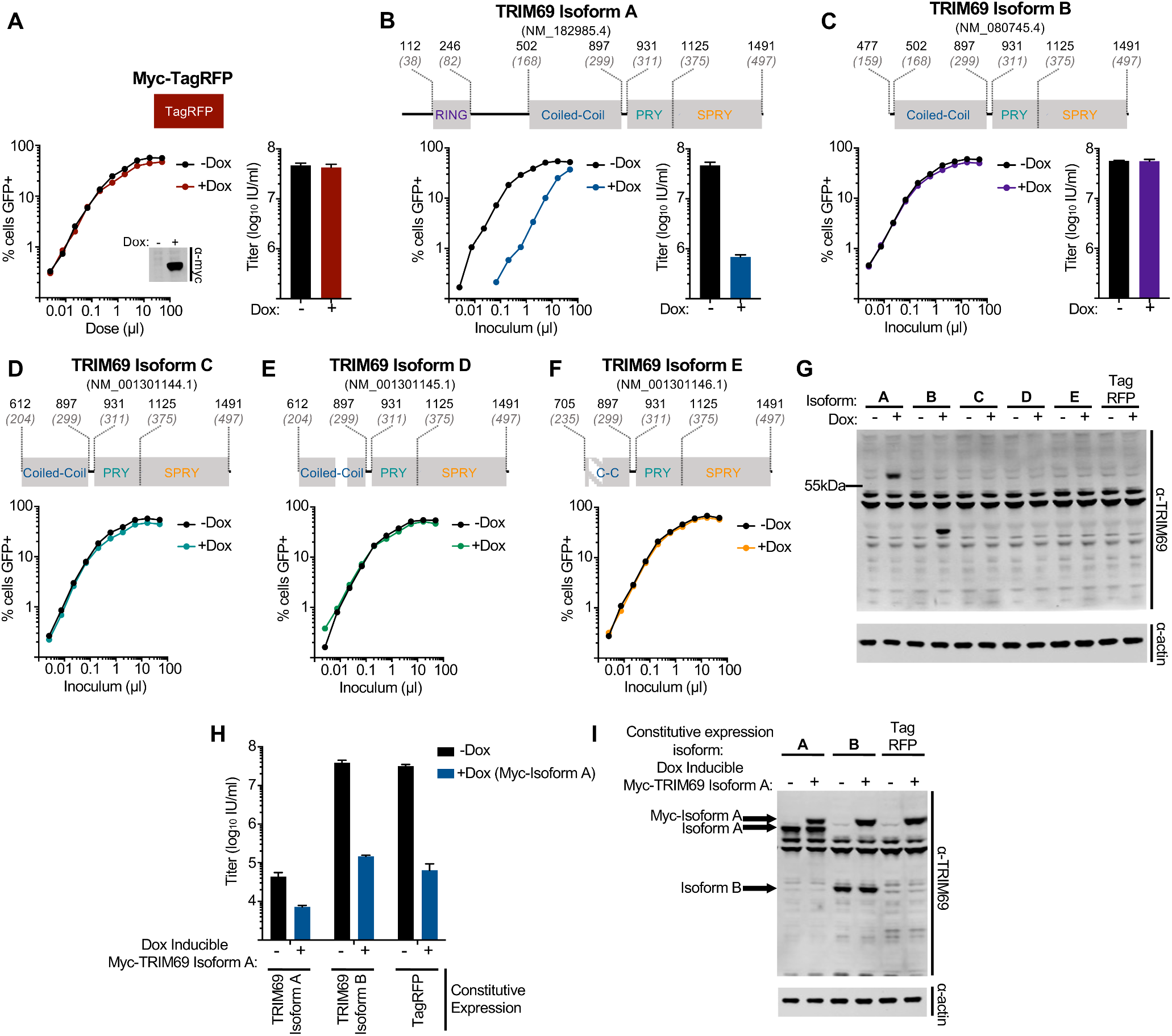
The anti-VSIV activity of TRIM69 appears limited to isoform A. (A) MT4 cells were modified to express myc-TagRFP in a doxycycline inducible fashion. A cartoon of TagRFP is depicted above typical titration curves of VSIV-GFP, the VSIV-GFP titres (16 h postinfection) and western blot (WB) analysis of Myc-TagRFP expression (24 h postinduction) in the presence and absence of doxycycline treatment. (B) MT4 cells were modified to express myc-TRIM69 isoform A in a doxycycline inducible fashion. A cartoon of TRIM69 isoform A is depicted above typical titration curves of VSIV-GFP alongside the VSIV-GFP titres of inducible TRIM69 isoform A expression in the presence and absence of doxycycline treatment. (C-F) As in B for (C) isoform B, (D) isoform C, (E) isoform D, and (F) isoform E. (G) WB analysis of TRIM69 expression in panels C-F. (H) Cells from panel B were modified to constitutively express TRIM69 isoform A, isoform B or TagRFP (by LHCX transduction). The titre of VSIV-GFP was determined in the presence and absence of doxycline inducible Myc-TRIM69 isoform A. (I) WB analysis of the cells in H. In all cases, virus titrations were carried out on at least 2 occasions and typical results are shown. Mean titers and standard deviation are plotted based on at least 3 doses (in the linear range).

As well as lacking antiviral activity, shorter isoforms of TRIM5 can act as dominant negative inhibitors of endogenous and exogenous TRIM5α-mediated restriction (34). We thus examined whether TRIM69 isoform B might similarly inhibit the anti-VSIV activity of isoform A. When TRIM69 (isoform A) was constitutively expressed, it conferred potent protection from VSIV-GFP infection (Figure 3H) and this protection was further enhanced by the presence of inducible myc-tagged TRIM69 (Figure 3H). In contrast, constitutively expressed TRIM69 isoform B had negligible effect on the ability of isoform A to inhibit VSIV (Figure 3H), despite being abundantly expressed (Figure 3I). Thus, although TRIM69 isoform B has no detectable antiviral activity, it does not appear to interfere with the ability of isoform A to block VSIV.

#### Multiple TRIM69 alleles circulate in human populations and TRIM69 exhibits strong signatures of positive selection

We next analysed structural variation at the TRIM69 locus in human populations, using data from The 1000 Genomes Project collated by ENSEMBL (35). Interestingly, multiple TRIM69 alleles circulate at high frequencies in human populations. Surprizingly, the human RefSeq (NM_182985) is only the third most common human allele, present at frequencies of ~4% (European) to ~17% (South Asian) in human populations. In light of this, we examined the ability of all the major TRIM69 alleles (with a frequency >5% in at least one population) to inhibit a small panel of vesiculoviruses (Figure 4). We cloned the seven main alleles into our doxycycline inducible expression system (Figure 4A-C) and challenged these cells with a small panel of vesiculoviruses (Figure 4C). In spite of the amino acid variation, all seven major alleles conferred potent protection from VSIV infection (Figure 4E). Although some variation in the magnitude of protection was observed, we attributed this to slight variations in TRIM69 expression levels, as opposed to variation in the anti-VSIV activity of the different alleles (Figure 4C). TRIM69 again exhibited exquisite antiviral specificity and none of the TRIM69 alleles inhibited VSV New Jersey (VSNJV) or chandipura virus (CHNV-GFP) (Figure 4F,G).

**Figure 4.**
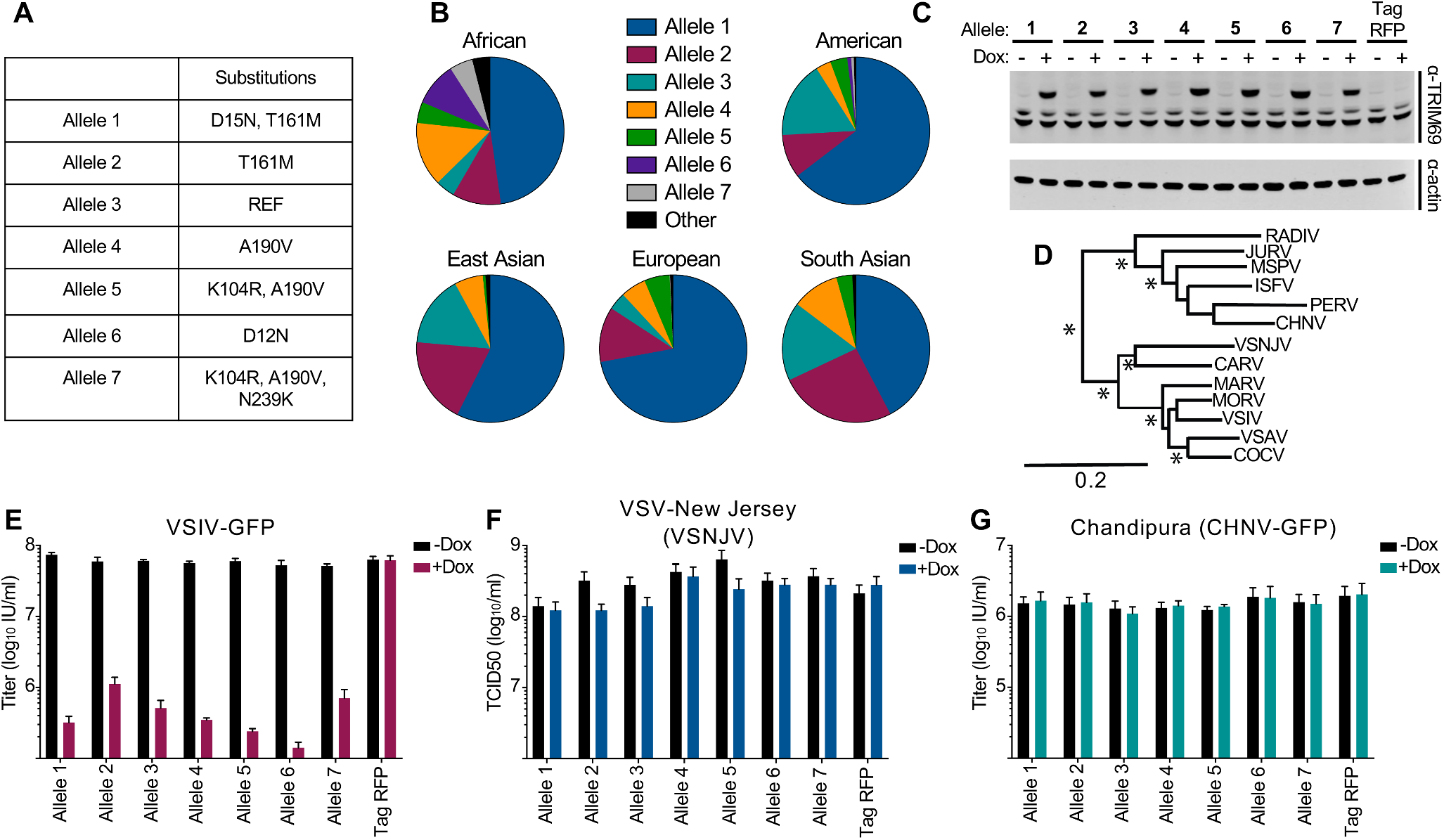
Multiple TRIM69 alleles circulate in human populations. (A) The amino acid substitutions of the most common TRIM69 alleles. (B) The allelic frequency of the major alleles by geographic region. (C) MT4 cells were modified to express the 7 most common TRIM69 alleles in a doxycycline inducible fashion. WB analysis of myc-TRIM69 expression in cells with and without 24 h of doxycycline treatment. (D) The redrawn phylogeny of common vesiculoviruses (75) is shown (asterisks represent bootstrap proportion higher than 85%). The cells from C were used to examine the ability of divergent TRIM69 alleles to inhibit VSIV-GFP (E), VSNJV (F) and CHNV-GFP (G). Virus titrations were carried out on at least 2 occasions and typical results are shown. Mean titers and standard deviation are plotted based on at least 3 doses (in the linear range).

Because multiple TRIM69 alleles currently circulate in human populations and antiviral TRIM proteins can possess strong signatures of positive selection (36), we conducted positive selection analysis of TRIM69 sequences from primates. We retrieved and aligned the TRIM69 coding region from 18 primate species representing >40 million years of divergent evolutionary pressures (37). Using the maximum likelihood approach in PAML (38), we tested whether models that permit positive selection on individual codons (dN/dS>1) were a better fit to these data, than models that do not allow positive selection. In each case, permitting sites to evolve under positive selection gave a better fit (Figure 5AB), with a high proportion of codons exhibiting dN/dS ratios greater than 1 (32.8% with average dN/dS=1.9). These analyses identified six residues exhibiting relatively strong signatures of positive selection (Figure 5AB). One of the sites identified using this approach (S404) is within the SPRY domain (the domain that forms the host-pathogen interface that defines the antiretroviral specificity of TRIM5 (36)). These analyses suggest that whilst the majority of sites (67.2%) in TRIM69 have evolved under purifying selection (in order to maintain the overall structure and function of TRIM69), positive selection has likely occurred at specific sites perhaps influencing the antiviral activity of TRIM69 (36).

**Figure 5.**
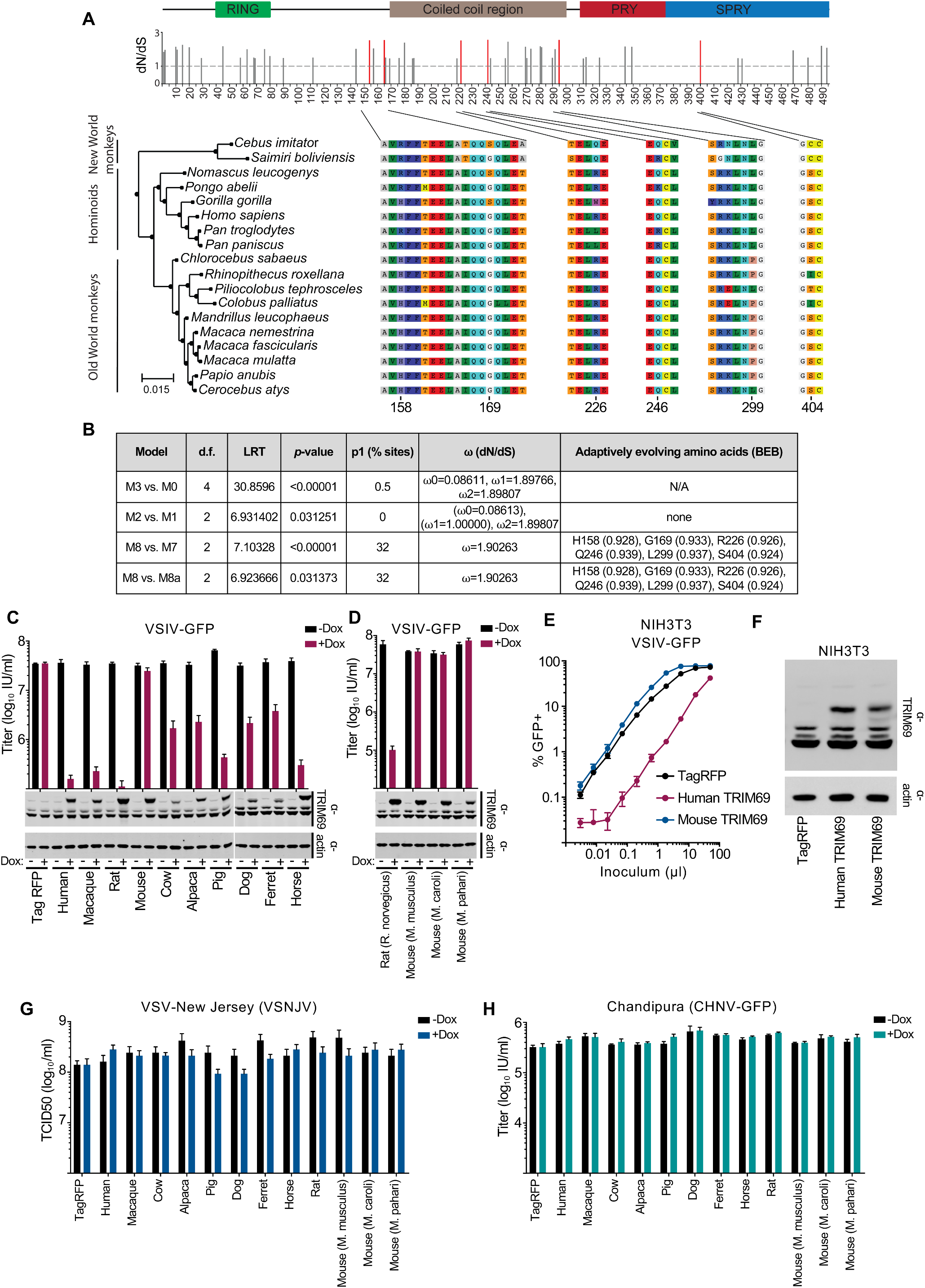
TRIM69 exhibits signatures of positive selection and anti-VSIV activity has been lost in the Mus lineage. (A) The diagram (above) illustrates the conserved domains of TRIM69 based on CDD v3.16 from NCBI. The bar chart (below) represents the dN/dS values of the sites with positive selection. The 6 sites with statistically significant positive selection (PAML M8, BEB, *P* > 0.9) are shown in red. The phylogenetic relationship between the 18 sequences spanning the hominoids is shown on the left with the regions of the alignment with statistically significant positive sites shown numbered on the right. (B) The results of the Chi-squared test comparing the PAML models is shown in the table along with the proportion of sites under positive selection and the average dN/dS for those sites. (C) MT4 cells were modified to express myc-TRIM69 orthologs from multiple species in a doxycycline inducible fashion. WB analysis of TRIM69 expression in cells with and without 24 h of doxycycline treatment is shown beneath the titres of VSIV-GFP (16 h postinfection) in the presence and absence of doxycycline treatment. (D) As in C, examining TRIM69 from rat and divergent mice. (E) NIH3T3 cells were modified to stably express human and mouse TRIM69 (LHCX) before being infected with serially diluted VSIV-GFP. (F) WB analysis of TRIM69 expression in the cells from E. The cells from C were used to determine the titers of (G) VSNJV and (H) CHNV-GFP. In all cases, virus titrations were carried out on at least 2 occasions and typical results are shown. Mean and standard deviation are plotted and titres are based on at least 3 doses (in the linear range).

Based on the signatures of positive selection at the TRIM69 locus in primates, we hypothesized that divergent TRIM69 proteins might exhibit divergent antiviral specificities. We therefore cloned a variety of TRIM69 orthologs from a selection of species (including primates and a variety of other mammals) into our doxycycline inducible expression system. Similar to human TRIM69, orthologs from rhesus macaques, rats, cows, alpacas, dogs, ferrets and horses all potently inhibited VSIV infection (Figure 5C). The magnitude of inhibition was variable, but we attributed the majority of this variability to different TRIM69 expression levels (Figure 5C). Strikingly, murine TRIM69 did not inhibit VSIV, despite being expressed at higher levels than multiple inhibitory orthologs (Figure 5C). Furthermore, the anti-VSIV activity of TRIM69 has apparently been lost in the Mus genus as TRIM69 orthologs from *M. caroli* and *M. pahari* were non-inhibitory, whereas rat TRIM69 potently inhibited VSIV (Figure 5D). This is not simply due to the murine orthologs lacking specific cofactors/interactions within human cells, as stable expression of murine TRIM69 (*M. musculus*) in mouse cells indicated that murine TRIM69 still possessed no apparent anti-VSIV activity (Figure 5E). Crucially, murine TRIM69 was abundantly expressed in these cells and human TRIM69 conferred potent protection from VSIV infection, when expressed at similar levels in the identical murine background (Figure 5EF). Thus, while murine cells support TRIM69-mediated anti-VSIV activity, murine orthologs of TRIM69 have lost the ability to inhibit VSIV.

We also challenged the species variants of TRIM69 with related vesiculoviruses (VSNJV and CHNV), but none of the orthologs considered possessed substantial antiviral activity against these viruses (Figure 5GH).

#### The VSIV phosphoprotein confers sensitivity/resistance to TRIM69

The short generation times inherent to the lifecycles of most viruses means that resistance to inhibition can be rapidly selected *in vitro* (5, 26, 39, 40). These in *vitro* evolution approaches can rapidly identify antiviral sensitivity/resistance determinants in viruses targeted by antiviral factors (39, 41). We used a diverse swarm of replication competent full-length VSIV-GFP (FL VSIV-GFP) (42) that had been propagated in mammalian cells. This parental virus stock was potently inhibited by TRIM69 following overnight infection (Figure 6A). We used this virus to inoculate a culture expressing TRIM69, using a dose where <1% of cells were GFP-positive following overnight incubation. Four days later, VSIV had begun to overwhelm this culture and the supernatant was filtered and transferred to a new culture (also induced to express TRIM69). In this second passage, VSIV replicated far more efficiently in the presence of TRIM69, suggesting resistance had emerged. Such rapid acquisition of resistance is a documented property of VSIV populations and VSIV fitness has been previously shown to increase >1000-fold in a single passage (within a new cellular environment) (43). We titrated the filtered supernatant containing the TRIM69-passaged swarm, in the presence and absence of TRIM69, and observed that VSIV had been selected to resist TRIM69 inhibition (Figure 6A). We sequenced the viral population and observed four substitutions that had been selected to near uniformity in the viral swarm (Figure 6BC). These substitutions included two synonymous changes and two nonsynonymous substitutions. Because we had previously observed that the VSV-G protein was not targeted by TRIM69 when used to pseudotype HIV-1 (Figure 1E), we reasoned that one substitution, E92K in VSV-G, was unlikely to confer resistance to TRIM69. This left only one non-synonymous substitution, D70Y, located in the VSIV phosphoprotein (P-protein) that might confer TRIM69-resistance. We mutated this residue in isolation, in the FL VSIV-GFP plasmid background and rescued the parental and D70Y mutant viruses. While the rescued parental virus was potently inhibited by TRIM69, the P-protein D70Y mutant was completely insensitive to inhibition by TRIM69 (Figure 6D). Thus, the VSIV P protein is the genetic target of TRIM69 and can determine sensitivity or resistance to TRIM69. Moreover, just a single amino acid within the P protein can determine TRIM69 sensitivity.

**Figure 6.**
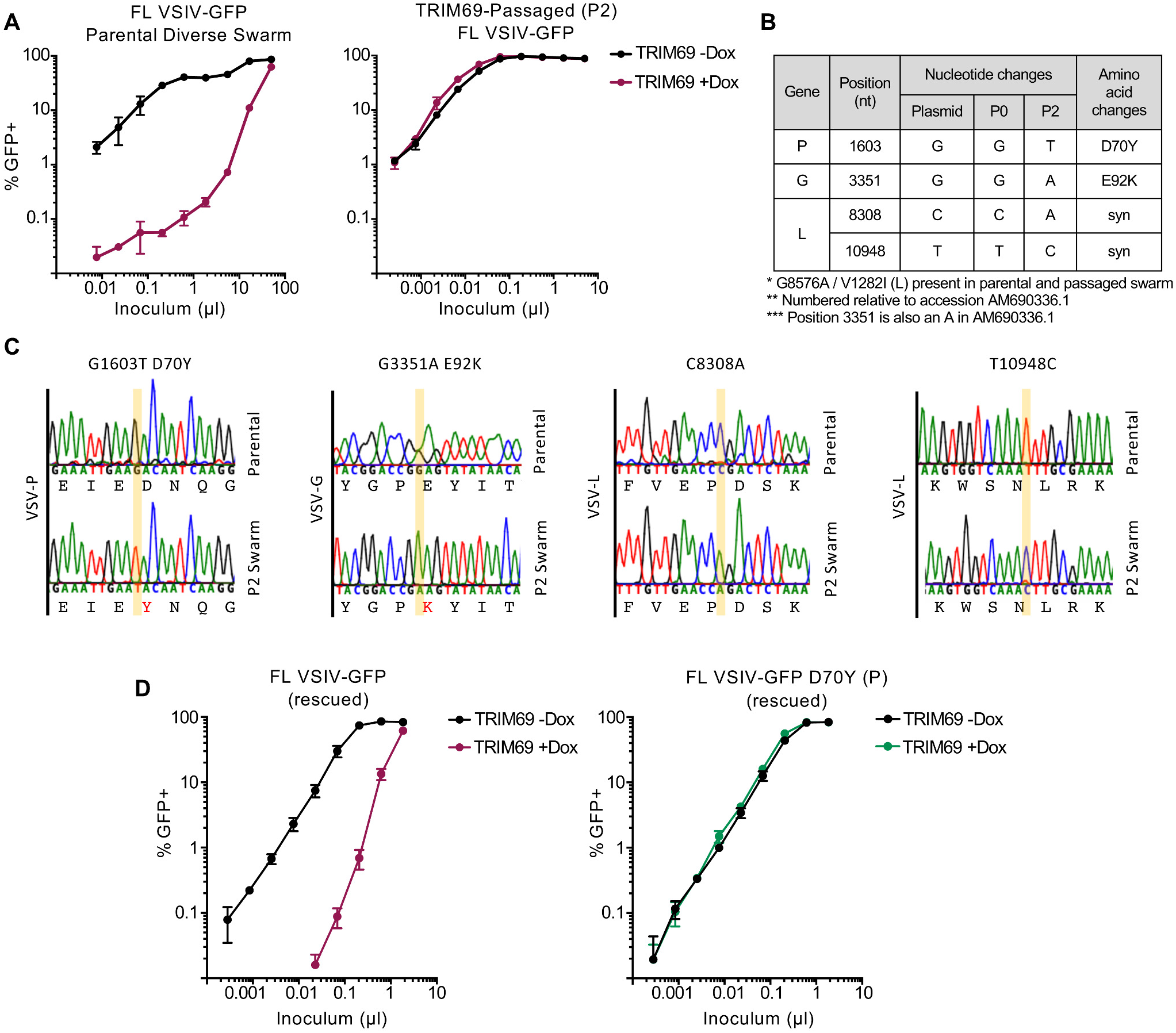
The VSIV phosphoprotein is a genetic susceptibility/resistance determinant of TRIM69 anti-VSIV activity. Serially diluted infection of MT4 cells modified to express doxycycline inducible human TRIM69 with (A) parental FL-VSIV-GFP or (B) FL-VSIV-GFP passaged twice in the presence of human TRIM69 is shown. (B) The passaged and parental swarms were sequenced and the listed changes were selected to near uniformity in the passaged population. (C) Sequencing chromatograms of directly-sequenced PCR products amplified from the reverse transcribed viral swarms are shown. (D) Titrated infection of the cells in A with the parental virus and the D70Y (P) mutant FL-VSIV-GFP both rescued in parallel. In all cases, virus titrations were carried out on at least 2 occasions and typical results are shown. Mean and standard deviation are plotted.

#### TRIM69 inhibits VSIV but not DENV2

During the preparation of this manuscript, it was reported that TRIM69 is an ISG that targets dengue virus type 2 (DENV-2), through an interaction with NS3, that targets NS3 for degradation (32). This TRIM69-mediated degradation of NS3 interrupts the lifecycle of DENV-2 (32). This observation was of immediate interest to us as it is unusual for an antiviral factor to specifically target viruses as divergent as DENV-2 (positive sense ssRNA virus) and VSIV (negative sense ssRNA virus). Moreover, murine TRIM69, which is inactive against VSIV, was reported to inhibit DENV-2 (32), suggesting species variants of TRIM69 might have divergent antiviral specificities. We thus compared the ability of TRIM69 to inhibit DENV-2 and VSIV in the same experiment. Because Vero cells are susceptible and permissive to both DENV-2 and VSIV, we selected these cells as the background for our experiments. We generated Vero cells that stably expressed human, rat or mouse TRIM69 and infected these cells with a titrated challenge of DENV-2 or VSIV-GFP. We used the New Guinea C strain of DENV-2 as this strain was previously reported to be inhibited by TRIM69 (32). Unexpectedly, DENV-2 produced in mammalian or insect cells was not inhibited by TRIM69 under these conditions (Figure 7 AD). In contrast, VSIV was potently inhibited by human and rat TRIM69 (Figure 7 EF) in parallel experiments.

**Figure 7.**
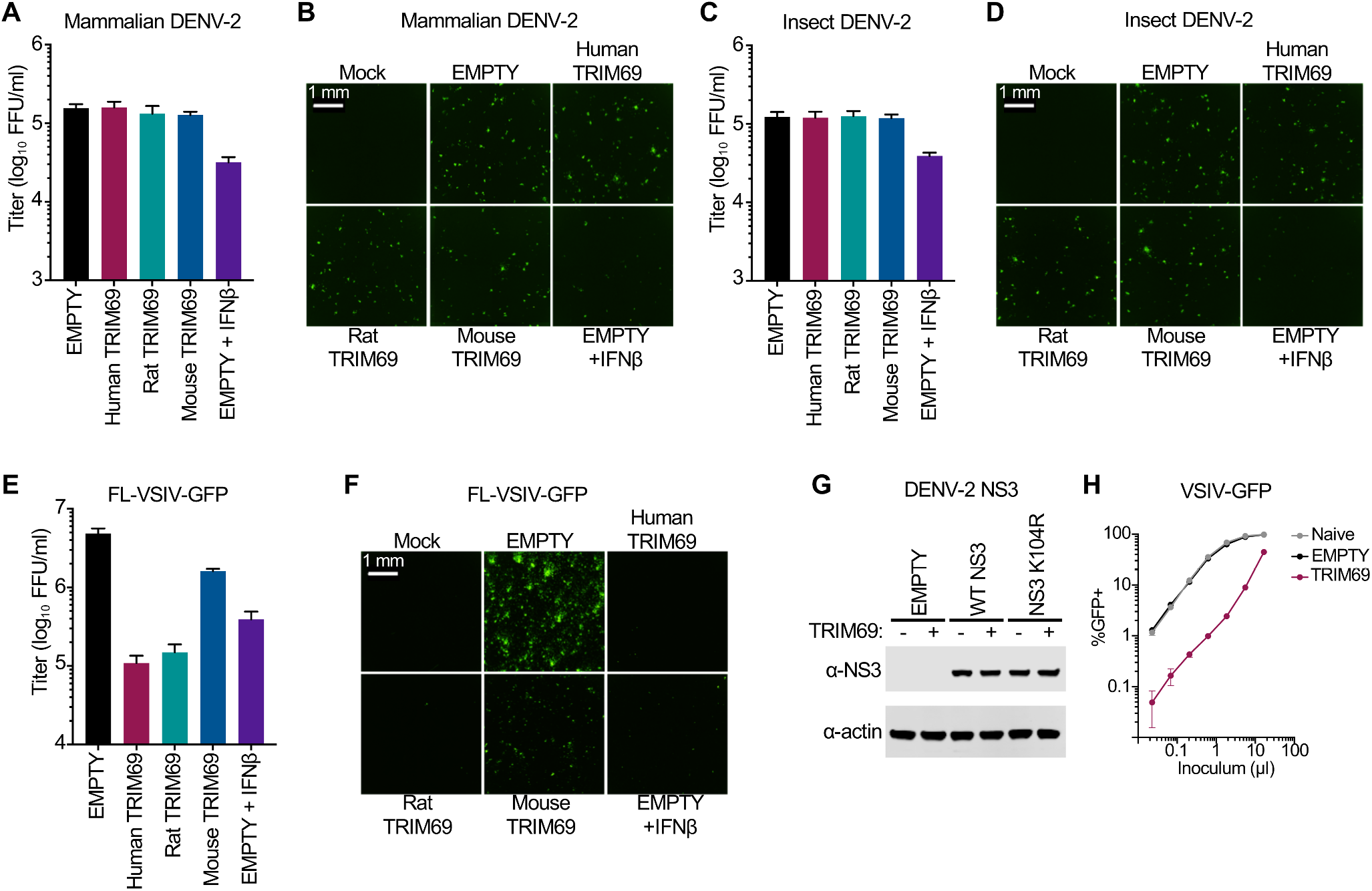
TRIM69-mediated inhibition of DENV-2 was not detected under conditions that restrict VSIV. Vero cells were modified to stably express human, rat and mouse TRIM69 or were transduced with the corresponding empty vector (SCRPSY). (A and B) Cells were challenged with DENV-2 propagated in Vero cells (Mammalian DENV-2) or (C and D) with DENV-2 produced from *Aedes albopictus* C6/36 cells (Insect DENV-2). 48 h after infection, cells were fixed and stained for DENV-2 infection and infected foci were imaged and enumerated using a Celigo imaging cytometer. (E and F) As in A-D, Vero cells were infected with FL-VSIV-GFP, fixed and analysed 16 h after infection as in A-D. (G) HEK 293T cells were modified to stably express human TRIM69 or were transduced with the corresponding empty vector (SCRPSY). Empty (-) and TRIM69-expressing (+) cells were transfected with DENV-2 NS3 expression plasmids and NS3 expression was analysed 48 h later using WB. (H) In parallel with the experiment in G, functional TRIM69 expression in equivalent HEK29T cells was assessed by titrated infection with VSIV-GFP. In all cases, virus titrations were carried out on at least 2 occasions and typical results are shown. Mean and standard deviation are plotted.

Because the previously published work had not specifically investigated exogenous TRIM69 expression in Vero cells, we examined whether TRIM69 expression in HEK 293T cells would induce the degradation of transfected NS3 but not NS3 K104R (recapitulating published observations) (32). Surprisingly, TRIM69 had no effect on NS3 expression levels (Figure 7G). In contrast, VSIV was potently blocked in parallel experiments using the equivalent cells (Figure 7H). We conclude that DENV-2 is not always inhibited by TRIM69, even under conditions where TRIM69 exhibits substantial antiviral activity against VSIV.

#### The anti-VSIV activity of TRIM69 is not dependent upon IFN-signalling or E3 ubiquitin ligase activity

TRIM69 has a RING domain predicted to have E3 ubiquitin ligase activity. Moreover, proteasomal degradation has previously been reported to be involved in the TRIM69-mediated inhibition of DENV-2 (32). In addition, TRIM69 isoform B (which lacks a RING domain) possess no anti-VSIV activity (Figure 3C). We therefore considered whether inhibiting proteasomal degradation might affect TRIM69 antiviral activity. As a less toxic alternative to MG132, we used bortezomib (Bort), an inhibitor of the 26S proteasome, which is also licensed for clinical use (44, 45). In order to validate the efficacy of inhibition, we transduced cells with a lentiviral vector encoding ubiquitin fused to GFP. This fusion protein is rapidly degraded and GFP expression is not visible under normal culture conditions (Figure 8AB). However, when the 26S proteasome was inhibited by bortezomib treatment, abundant GFP expression was visible, suggesting proteasomal inhibition was efficient in our culture system. In parallel experiments, we considered the ability of proteasome inhibition to influence TRIM69-mediated restriction of VSIV. When normalised to infection in the absence of TRIM69, proteasomal inhibition appeared to partially rescue the restriction of VSIV (Figure 8C). However, bortezomib treatment caused noticeable toxicity in these experiments, consistent with bortezomib’s pro-apoptotic and anti tumor cell growth properties (45). Visual inspection of the titration curves indicated that the majority of the rescue was due to reduced VSIV infection in control cells (in the presence of bortezomib), as opposed to proteasomal inhibition actually enhancing infection in the presence of TRIM69 (Figure 8D). Importantly, TRIM69 potently restricted VSIV (>50-fold), in the presence of efficient proteasome inhibition, indicating that proteasomal degradation is not necessary for effective TRIM69-mediated restriction.

**Figure 8.**
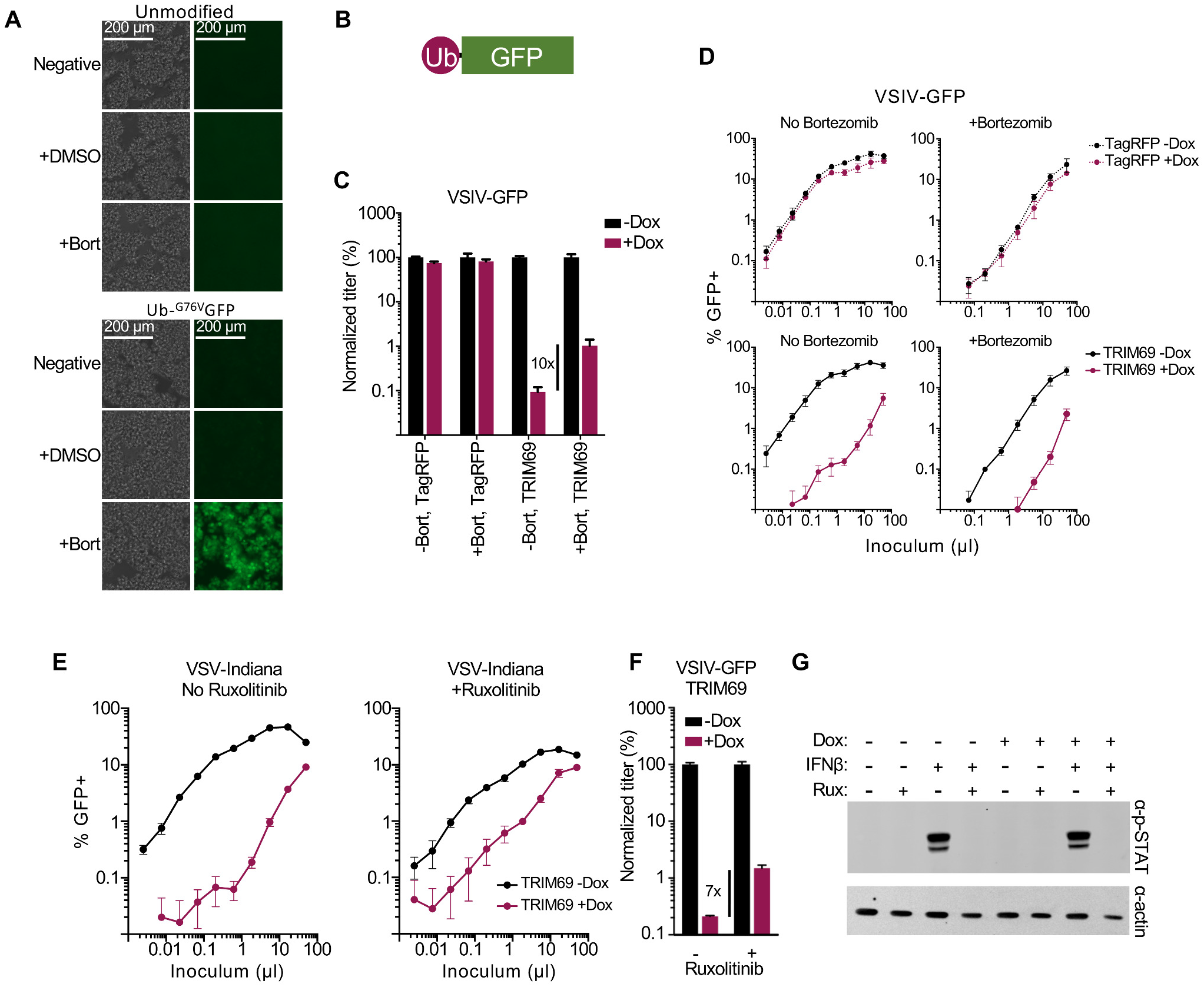
TRIM69-mediated antiviral activity does not require the proteasome or IFN-signalling. (A) phase contrast and epifluorescence images (EVOS FL microscope) of unmodified MT4 cells or MT4 cells modified to express Ub-^G76V^GFP in the presence or absence of 50 nM bortezomib (proteasome inhibitor). (B) a cartoon of Ub-^G76V^GFP. (C and D) MT4 cells modified to express inducible myc-TagRFP or myc-TRIM69 (human) were treated or not treated with doxycycline (24 h) before being infected with VSIV-GFP (16 h) in the presence or absence of 50 nM bortezomib. Normalized titers are shown in C of the titration curves in D. (E) As in C and D examining VSIV-GFP in the presence of inducible myc-TRIM69 in the presence and absence of 2 μM ruxolitinib (JAK1/2 inhibitor). Normalized titers are shown in F of the titration curves in E (G) Phospho-Stat1 expression in in the presence and absence of 2 μM ruxolitinib incubated for 24 h in the presence or absence of 1000 units of IFNβ examined in MT4 cells measured in parallel with the cells in E. In all cases, virus titrations were carried out on at least 2 occasions and typical results are shown. Mean and standard deviation are plotted.

Many TRIM proteins are involved in antiviral signalling and appear to have antiviral activity because their exogenous expression promotes or potentiates a polygenic antiviral state (29, 30). Although the observation that most viruses escape inhibition by TRIM69 suggests that a polygenic response is not involved, we investigated whether TRIM69 could inhibit VSIV in the absence of JAK-STAT signalling. We therefore used the janus kinase inhibitor ruxolitinib to block interferon signalling in TRIM69 expressing cells (46). We then examined the ability of TRIM69 to block VSIV infection in the absence of IFN-signalling. As with bortezomib, some toxicity was observed and ruxolitinib slightly reduced the level of infection in the absence of TRIM69 (Figure 8E,F). Thus, the majority of the apparent ~7-fold rescue (Figure 8F) was caused by reduced infection in the absence of TRIM69 (as opposed to ruxolitinib solely enhancing infection in the presence of TRIM69). However, in contrast to bortezomib, ruxolitinib did modestly increase the amount of VSIV infection in the presence of TRIM69 (increasing the titer ~2.5-fold) (Figure 8E). Importantly, the ruxolitinib treatment effectively blocked IFN-signalling and no phosphorylated STAT1 was detectable following IFN-treatment in the presence of ruxolitinib (Figure 8G). Notably, TRIM69 expression alone was insufficient to trigger the phosphorylation of STAT1 (Figure 8G) and TRIM69 potently blocked VSIV (>20-fold) in the presence of ruxolitinib. Thus, IFN-signalling is not necessary for TRIM69 to inhibit VSIV (although JAK-STAT signalling might modestly enhance this inhibition).

## DISCUSSION

Although vesiculoviruses can be particularly sensitive to IFN-induced inhibition, the specific ISGs that mediate the anti-VSIV activity of IFNs (and how this inhibition is achieved) have not yet been fully defined. Here we show that one ISG, TRIM69, can potently and specifically inhibit VSIV. Moreover, the high degree of specificity of this inhibition overwhelmingly suggests that this inhibition is direct. TRIM69 blocked VSIV whereas other vesiculoviruses (and more divergent RNA viruses) were entirely resistant. Furthermore, the inhibition of VSIV appeared to be remarkably specific, as a single substitution conferred complete resistance to TRIM69. Thus, although we cannot exclude the possibility that TRIM69 expression elicits some kind of antiviral signalling, such a signalling event would have to be independent of proteasomal degradation and JAK-STAT signalling, and also result in an outcome that specifically targeted VSIV. Therefore, the more parsimonious explanation is that TRIM69, or a complex containing TRIM69, directly targets VSIV (and interferes with the virus lifecycle). This potentially adds TRIM69 to an expanding list of directly antiviral TRIM proteins that includes the classical restriction factors TRIM5 (34) and PML (47). Nonetheless, inhibition of IFN signalling modestly rescued VSIV infection and TRIM69 KO slightly reduced the IFN-induced inhibition of HIV-1. Thus, it is possible that TRIM69 might also play a signalling role, analogous to the signalling capacity of established directly-antiviral factors such as TRIM5, TRIM21 and tetherin (48–50).

The most surprizing aspect of the highly specific anti-VSIV activity of TRIM69 was that DENV-2, previously reported to be inhibited by TRIM69 (32), was not inhibited in our experiments. Importantly, this lack of TRIM69-sensitivity was observed whilst parallel cultures (of the equivalent cells) fiercely resisted VSIV infection. There are many possible explanations for these apparently contradictory observations. The most likely explanation is that due to the highly specific nature of TRIM69-mediated inhibition, some aspect of the virus strain, cellular background or method used has led to DENV-2 appearing insensitive to inhibition (in our experiments). We were careful to use the same strain as used previously (specifically the New Guinea C strain, which we obtained from Public Health England). We confirmed the presence of the ‘TRIM69-sensitive’ lysine (at position 104 of NS3) in our virus stocks and more work will be required to understand why we did not observe DENV-2 inhibition in the face of antiviral TRIM69. Importantly, with both VSIV and DENV-2, a single amino acid substitution conferred resistance to TRIM69. Thus, seemingly minor differences in viral strains could easily reconcile these apparently contradictory observations.

Although we do not know how TRIM69 impedes VSIV, the block must occur relatively early in the viral lifecycle. TRIM69 is able to block VSIV prior to the expression of GFP, which in the case of the rVSVΔG-GFP system is encoded in place of the VSV-G protein. Thus, the TRIM69-mediated block must occur at a step prior to translation of VSV-G subgenomic RNA. In the absence of direct evidence, the D70Y mutation in the VSIV phosphoprotein does provide some potential mechanistic clues. Importantly, D70 is present in a region of VSV-P that is heavily phosphorylated (phosphorylation of residues 60, 62 and 64 has been described) (51). Phosphorylation of these sites has been proposed to be important in P-protein dimerization and interaction with the L-protein (large polymerase protein) (52–56) and phosphorylation of this region is essential for functional RNA dependent RNA polymerase activity (57). Because tyrosine residues are known to be phosphorylated in the VSV-P protein (58), it is therefore tempting to speculate that a tyrosine at position 70 could be phosphorylated, and that this phosphorylation somehow overcomes the block mediated by TRIM69. A speculative mechanism that involves TRIM69-mediated inhibition of VSIV RNA dependent RNA polymerase activity (by inhibiting the association of P and L-proteins) would be entirely consistent with the early block described herein.

The N-terminal region of vesiculoviral P-proteins are relatively divergent, providing a possible explanation for the observed insensitivity of VSNJV and CHNV to TRIM69. Thus, examining the more closely related Morreton virus (MORV), Vesicular Stomatitis Alagoas Virus (VSAV), Cocal virus (COCV) or Maraba viruses (MARV) might identify more viruses which are sensitive to TRIM69, and potentially identify further evidence of species specificity. Notably, after analysing 49 VSIV sequences (deposited in GenBank), we did not observe a tyrosine at position 70 of VSIV-P in any VSIV sequences. This suggests that if TRIM69 substantially inhibits VSIV *in vivo*, the D70Y mutation must be deleterious for some other reason and is therefore negatively selected (despite conferring resistance to TRIM69).

Our failure to identify any viruses other than VSIV that were inhibited by TRIM69, limited our ability to identify whether species variants of TRIM69 possess divergent antiviral specificities. Despite this, differential activity was observed as anti-VSIV activity has apparently been lost in the Mus genus. Whether murine TRIM69s have been selected to target other viruses or whether murine orthologs have no antiviral activity at all remains to be determined. Thus, murine TRIM69 might be an active antiviral factor selected to target a divergent spectrum of viruses, in a way that prevents it from inhibiting VSIV. The rat and mouse TRIM69 variants were the closest orthologs we tested, that displayed differential activity against VSIV, and these species variants differ at 38 amino acid positions. Despite detecting site-specific signatures of positive selection in TRIM69, all of the signature sites are conserved between rats and mice. Thus, the genetic basis of the differential anti-VSIV activity observed in rodent TRIM69 variants is currently unknown.

The lack of anti-VSIV activity conferred by murine TRIM69 could have important implications for our understanding of how the IFN response constrains VSIV. Many key experiments have been conducted in mice (15, 16) and it will be important to establish, in future, whether non-murine variants of TRIM69 possess anti-VSIV activity *in vivo*, potentially limiting the ability of VSIV to invade and colonise certain tissues. It is possible that the potent inhibition of VSIV observed *in vitro* will be recapitulated *in vivo*. In this way, IFN-induced TRIM69 might inhibit natural VSIV infections and similarly influence the safety and/or efficacy of therapeutic interventions (based upon VSIV). Thus, an improved understanding of how TRIM69 inhibits VSIV may help us to better understand VSIV pathogenesis and eventually lead to tangible benefits in the clinical use of VSIV derivatives.

## MATERIALS AND METHODS

### Cells and viruses

Adherent HEK 293T, BHK21, BSRT7/5 cells (modified to stably express T7 RNA polymerase (59)), NIH 3T3 and Vero cells were propagated from lab stocks maintained in DMEM supplemented with 9% fetal calf serum (FCS) and 20 μg/ml gentamicin. CADO-ES1 semi-adherent cells were purchased from the DSMZ (ACC 255) and suspension MT4 cells were expanded from lab stocks and maintained in RPMI supplemented with 9% FCS and 20 μg/ml gentamicin. C6/36 cells (*Aedes albopictus*) were propagated from existing lab stocks and maintained in L-15 (Leibovitz) media with GlutaMAX, 10% Fetal bovine serum, 10% tryptose phosphate broth and 1% penstrep. Transduced cells were selected and cultured in medium additionally supplemented with 2 μg/ml puromycin (Melford laboratories) or 200 μg/ml of hygromycin B (Invitrogen).

The VSIV-GFP virus (rVSVΔG-GFP) competent to undergo a single round of infection but not encoding the VSV-G envelope (rVSV-ΔG-GFP decorated with VSV-G expressed in trans) system was used (27). Virus stocks were generated as described previously (40). Briefly, HEK 293T cells were transfected with a VSV-G expression plasmid. The next day, the cells were infected with rVSV-ΔG-GFP using an MOI of 1. Progeny VLP stocks were harvested at 24 h postinfection and clarified using a 0.45 μm filter. The replication competent FL-VSIV-GFP, VSIV and VSNJV viruses were a generous gift from Megan Stanifer (Heidelberg University). Virus stocks were generated through infection of BHK21 cells using a low MOI. Once CPE was readily apparent, supernatants were harvested and clarified using a 0.45 μm filter.

The D70Y mutant (VSV-P) was made using the Agilent QuikChange Lightning Site-Directed Mutagenesis kit, in accordance with the manufacturer’s instructions, using the parental FL-rVSV-GFP plasmid as template (pVSV1(+)-GFP) (42) and the following oligonucleotides, 5’-GCT TCC GGA TCT GGT ACA TAC AAG CCT TGA TTG TAT TCA ATT TCT GGT TCA GAT TCT GT-3’ and 5’-TGA TTC TGA CAC AGA ATC TGA ACC AGA AAT TGA ATA CAA TCA AGG CTT GTA TGT ACC AG-3’. The entire coding region was subsequently sequence verified to confirm the sole presence of the D70Y (VSV-P) mutation. To rescue the parental and mutant virus, BSR-T7 cells were seeded and infected with FP-T7 virus (equivalent to MOI ~2 determined using DF-1 cells) (60). After a 1 h incubation, the FL-VSIV-GFP rescue plasmid (pVSV1(+)-GFP) or mutant D70Y (pVSV1(+)-GFP P-D70Y), were cotransfected together with pBS-N, pBS-P and pBS-L (KeraFAST) using FuGENE6 (Promega) with 1.66 μg, 0.83 μg, 0.50 μg and 0.33 μg respectively. After 48 h, GFP positive cells and CPE were observed in the transfected BSR-T7 cells. Supernatant containing VSIV was harvested and clarified using a 0.45-μM filter and stored at −80°C. VSIV was propagated in BHK-21 cells in T-25 flasks. Cells were infected with a low MOI of VSV harvested from BSR-T7 cells. At 48 h postinfection, (or when the majority of cells were GFP positive), supernatant was harvested and clarified using a 0.45-μM filter.

DENV2 was obtained from Public Health England (catalogue number 0006041v). The NS3 of the DENV2 stocks (propagated in Vero cells) used herein was directly sequenced. Briefly, RNA was extracted from infected cells using the same approaches described below for passaged FL-VSIV-GFP and the NS3 region was amplified using the following primers 5’-CGA AGA GGA AGA ACA AAT ACT GAC C-3’, 5’-GAT TGT ACG CCC TTC CAC CTG CTT C-3’ and PCR products sequenced using these primers and an additional primer (5’-GTG GAG CAT ATG TGA GTG CTA TAG C-3’). The NS3 sequence differed by one amino acid, a threonine at position 442, from GenBank: AAC59275 (strain New Guinea C).

The replication competent proviral clone NHG (JQ585717) and Δ*env* derivatives have been described previously (5, 61). Virus stocks were generated through transient transfection of HEK 293T cells in isolation (NHG) or in conjunction with a VSV-G expression plasmid (NHGΔ*env*). The single round RVFV system has also been described previously (62). Briefly, BHK-Rep cells were transiently transfected with pCAGGs-M. The cells were washed the following day and supernatant was harvested 48 h posttransfection and clarified using a 0.45-μM filter.

An NS1-eGFP expressing A/Puerto Rico/8/1934 (H1N1) virus was designed based on the previously described ‘Color-Flu’ system (63). A DNA sequence was synthesized (Genewiz) with flanking BsmBI sites corresponding to the NS segment of PR8 (GenBank accession EF467817.1) in which the NS1 ORF had been altered to code for a C-terminally eGFP-tagged NS1 protein with a CSGG linker. This was immediately followed in-frame by a CSG linker, the 2A protease of Porcine Teschovirus (PTV), and the NEP ORF. A splice acceptor site in the NS1 ORF was removed by introducing a527c and a530g (numbering in accordance with EF467817.1). The sequence was sub-cloned into the pHW2000 reverse genetics plasmid. The PR8-NS1-eGFP virus was rescued using a well-established reverse genetics system previously described (64) a generous gift from Ron Fouchier.

### Retroviral vectors and plasmids

The SCRPSY (KT368137.1) and doxycycline-inducible (LKOΔ-MycTagRFP-IP) lentiviral vectors have been previously described (5, 41). The retroviral vector LHSXN (a gift from T. Zang and P. Bieniasz) is a derivative of LHCX (Clontech) modified to contain the following MCS 5’-*AAG CTT* GGC CGA GAG GGC CGA AAA CGT TCG CGG CCG CGG CCT CTC TGG CCG *TTA AC-3’* between the *Hind*III and *Hpa*I sites (highlighted in italics) of LHCX. All human and species variants of TRIM69 were synthesized by Genewiz, based on NCBI sequences of the longest isoform, unless otherwise noted, from: human (human isoform A: NM_182985.4, isoform B: NM_080745.4, isoform C: NM_001301144.1, isoform D: NM_001301145.1, isoform E: NM_001301146.1), macaque (*M. mulatta:* XM_015142131.1), rat (*R. norvegicus:* BC091171 (ordered from Source Bioscience IRBPp993F0532D (IMAGE ID: 7132390))), mice (*M. musculus:* ordered from Source Bioscience IRAWp5000E114D (IMAGE ID: 6774293)), *M. caroli:* XM_021183811.1 and *M. pahari:* XM_021193471.1), cow (*B. indicus:* XM_019967990.1 (edited to change an ambiguity base at position 547 to a C to match the sequence of *B. taurus* XM_015473308)), alpaca (*V. pacos:* XM_015237089.1 (edited to remove 55 amino acids from the start of the sequence and replaced with ATGGAG (ME) that is found in all other TRIM69 species variants)), pig (*S. scrofa:* Ensembl ID: ENSSSCT00000005168 (edited to remove a 5’ K and replaced with ATGGAG [ME])), dog (*C. lupus:* XM_535459.6), ferret (*M. furo:* XM_004751292.2), horse (*E. caballus:* XM_014733851), and lizard (*A. carolinensis:* XM_008120705.1) sequences and cloned (using directional *SfiI* sites) into pSCRPSY, pLKOΔ-Myc-IP or pLHSXN as indicated in the text, figure or figure legend. Human allelic variants were cloned using overlap extension PCR and the following oligos: allele 1 5’-CTC TCT GGC CGA GAG GGC CAT GGA GGT ATC CAC CAA CCC CTC CTC CAA CAT CGA TCC AGG CAA CTA TGT TGA AAT GAA TGA TTC AAT C-3’, 3’-TGA CCC TGT TGG ATG GCA AGC TCC TCC ATG AAG AAA TGG ACA GCA TCA GAG ATT TGC AG-5’, 5’-GCA AAT CTC TGA TGC TGT CCA TTT CTT CAT GGA GGA GCT TGC CAT CCA ACA GGG TCA AC-3’, and 3’-TCT CTC GGC CAG AGA GGC CTT ACT GTG GAT GTA AGA TGT GCA ATG G-5’; allele 2 5’-CTC TCT GGC CGA GAG GGC CAT GGA GGT ATC CAC CAA CCC CTC CTC-3’, 3’-TGA CCC TGT TGG ATG GCA AGC TCC TCC ATG AAG AAA TGG ACA GCA TCA GAG ATT TGC AG-5’, 5’-GCA AAT CTC TGA TGC TGT CCA TTT CTT CAT GGA GGA GCT TGC CAT CCA ACA GGG TCA AC-3’, and 3’-TCT CTC GGC CAG AGA GGC CTT ACT GTG GAT GTA AGA TGT GCA ATG G-5’; allele 4 5’-CTC TCT GGC CGA GAG GGC CAT GGA GGT ATC CAC CAA CCC CTC CTC-3’, 3’-CAG ATG TAG CTT GTT TTC CTT GTG AGC AAC AAT AGC TTC CTT CTG CAT GTT CCT CAG GG-5’, 5’-CCT GAG GAA CAT GCA GAA GGA AGC TAT TGT TGC TCA CAA GGA AAA CAA GCT ACA TCT GC-3’, and 3’-TCT CTC GGC CAG AGA GGC CTT ACT GTG GAT GTA AGA TGT GCA ATG G-5’; allele 5 5’-CTC TCT GGC CGA GAG GGC CAT GGA GGT ATC CAC CAA CCC CTC CTC-3’, 3’-GTG GAT GGC CCT TGA GTA AGG GTA ACT TCC TAA TCT TCT CTA CCA ACT TGT CCA GTA CAG-5’ 5’-GTA CTG GAC AAG TTG GTA GAG AAG ATT AGG AAG TTA CCC TTA CTC AAG GGC CAT CCA CAG-3’, and 3’-TCT CTC GGC CAG AGA GGC CTT ACT GTG GAT GTA AGA TGT GCA ATG G-5’; allele 6 5’-CTC TCT GGC CGA GAG GGC CAT GGA GGT ATC CAC CAA CCC CTC CTC CAA CAT CAA TCC AGG CGA CTA TGT TGA AAT GAA TG-3’, and 3’-TCT CTC GGC CAG AGA GGC CTT ACT GTG GAT GTA AGA TGT GCA ATG G-5’; allele 7 5’-CTC TCT GGC CGA GAG GGC CAT GGA GGT ATC CAC CAA CCC CTC CTC-3’, 3’-GAG ACA TTG CTC CTG AAG CTG GCT CAG TTT CAA CTC CAT CTC CTC ATT CAA GGC TTT C-5’, 5’-GCC TTG AAT GAG GAG ATG GAG TTG AAA CTG AGC CAG CTT CAG GAG CAA TGT CTC TTA GC-3’, and 3’-TCT CTC GGC CAG AGA GGC CTT ACT GTG GAT GTA AGA TGT GCA ATG G-5’. Allele 4 was used as a template for Allele 5 PCRs and allele 5 was used as a template for allele 7. The N-terminal fusion of a mutated uncleavable ubiquitin moiety fused to GFP (Ub-^G76V^GFP) has been described previously (PMID: 10802622). Plasmid DNA containing Ub-^G76V^GFP, a gift from A. Fletcher (pcDNA3.1(+).Ub-^G76V^GFP), was digested with BamHI and *NotI* and inserted into the similarly digested lentiviral vector pCSGWΔ*Not*I (65) (a gift from G. Towers and A. Thrasher).

Gene editing was achieved using the lentiGuide-Puro system (66) in accordance with the ZhangLab protocols. The following oligos were used to make TRIM69 guides: 5’-CAC CGC AAC CCT GTA CTG GAC AAG T-3’, 5’-AAA CAC TTG TCC AGT ACA GGG TTG C-3’ (guide 1) and 5’-CAC CgA AGA AGT TAC CCT TAC TCA A-3’ and 5’-AAA CTT GAG TAA GGG TAA CTT CTT C-3’. Diploid CADOES1 cells were either transduced with vectors encoding Cas9 and the relevant TRIM69-targeting sgRNAs or transduced in parallel with vectors encoding Cas9 and no sgRNA (No guide). Single cell clones were generated using limiting dilution and the KO was confirmed by extracting genomic (DNeasy, Qiagen), followed by PCR amplification of the guide target regions in exon 2 using the following oligos 5’-CAC TTT CAA AGG AGA GAT TAT GTG C-3’ and 5’-GAG CAG TCT GGG CTT TCT AAT CAT C-3’. The PCR products were cloned into pGEM-T-Easy (Promega) and multiple clones were Sanger-sequenced.

Viral vectors were produced using transient transfection of HEK 293T cells (5 μg of vector/genome plasmid, 5 μg of the relevant GagPol expression vector and 1 μg of a VSV-G expression plasmid). Vector-containing supernatants were filtered (using a 0.45 μm filter) and were used to transduce the relevant cell types.

To make the NS3 expression plasmids, the NS3 coding sequence from DENV-2 New Guinea C strain (nucleotides 6376 – 6756 of GenBank accession KM204118.1), was synthesized (Genewiz), with a 5’ ATG, flanked by 5’ *Hind*III and 3’ *XbaI* sites and subcloned into pcDNA 3.1 (+). The K104R mutation (AAA -> AGA) was introduced using Agilent’s QuikChange Lightning Mutagenesis kit and the following primers: 5’-GAC GGC TCT TGG ATT TCT TCC AGG CTC CAA TGC-3’ and 5’-GCA TTG GAG CCT GGA AGA AAT CCA AGA GCC GTC-3’.

### Arrayed ISG expression screening

The ISG screens were executed as described previously (5, 26). Briefly, MT4 cells were seeded in 96-well plates and transduced with ISG-encoding SCRPSY vectors (one ISG per well). 48 h posttransduction, cells were infected with VSIV-GFP. Following incubation overnight, cells were fixed and analyzed using flow cytometry.

### Western blotting

For preparation of cell lysates, cell pellets were resuspended in SDS sample buffer (12.5% glycerol, 175 mM Tris-HCl [pH 8.5], 2.5% SDS, 70 mM 2-mercaptoethanol, 0.5% bromophenol blue). Proteins were subsequently separated on NuPage 4 to 12% Bis-Tris polyacrylamide gels and transferred onto nitrocellulose membranes. Blots were probed with either anti-actin (JLA20 hybridoma, courtesy of the Developmental Studies Hybridoma Bank, University of Iowa), one of two anti-TRIM69 antibodies (Abcam cat: ab111943 or Thermo Fisher cat: PA5-12215), anti-DENV2 NS3 (Thermo Fisher cat: PA5-32199), anti-phospho-STAT1 (Tyr701) (58D6), or anti-c-myc (9E10 hybridoma, Developmental Studies Hybridoma Bank, University of Iowa). Thereafter, membranes were probed with DyLight labeled goat secondary antibodies (Thermo) and scanned using a LiCor Odyssey scanner.

### Virus infections and titrations

For assays using GFP-encoding viruses, cells were seeded in 96 well plates. Adherent and semi-adherent cells were seeded 24 hours before challenge or treatment whereas suspension cells were seeded immediately prior to infection or treatment. In experiments using doxycycline inducible TRIM69 expression, cells were treated with 125 ng/ml of doxycycline hyclate (Sigma) 24 h before infection. Where stated, cells were treated with 2 μM ruxolitinib (INCB018424, Stratech, S1378-SEL) or 50 nM bortezomib (Cell Signaling Technology #2204) or 1000 units of IFNβ (pbl Assay Science, cat: 11420-1) immediately before infection (or in the case of IFNβ, 24 hours before infection, or 4 hours treatment for the data in Figure 7). Cells were then infected with titrated challenges of the indicated virus and incubated overnight (~16 h) or for 48 h (HIV-1 and RVFV) prior to fixation using 4% formaldehyde and enumeration of infected GFP-positive cells using flow cytometry.

For quantification of focus forming units (FFUs), Vero cells were seeded in 96-well plates. The following day, cells were pretreated with IFNβ (24h at 1000U/ml) prior to infection with FL-VSIVGFP or DENV-2 propagated in Vero cells (Mammalian) or C6/36 cells (Insect). Cells were infected with titrated DENV-2 in DMEM (2% FCS) for one hour prior to overlay with DMEM (5% FCS, 0.8% carboxymethylcellulose). 48 h later, cells were fixed (methanol) and permeabilized (0.1 % Triton X-100, Fisher). FFUs were visualized using MAB8705 Anti-dengue Virus Complex Antibody clone D3-2H2-9-21 (Millipore) and goat anti-mouse Alexa Fluor 488 (A-11001, Thermo Fisher) as described previously (67). As a control, serially diluted FL-VSIV-GFP was examined in parallel and fixed in 4% formaldehyde following overnight incubation (~16 h). The number of fluorescent foci of immunostained DENV-2 and VSIV infected cells was enumerated using a Celigo imaging cytometer (Nexcelom, Bioscience).

For TCID50 assays, MT4 cells were seeded in 96-well plates and treated with 125 ng/ml of doxycycline hyclate (Sigma) 24 h prior to infection. The cells were infected with 8 replicates of 3-fold serially diluted full-length VSNJV or VSIV. After 72 hpi, CPE was analysed and the TCID50 was calculated using the Spearman & Kärber algorithm using a modified TCID50 calculator from Marco Binder.

### DENV-2 NS3 transfections

HEK 293T cells were transduced with either SCRPSY-EMPTY or SCRPSY-TRIM69 and seeded in 6-well plates before transfection with 2 μg of pcDNA (either EMPTY, wild type DENV2 NS3, or DENV2 NS3 K104R mutant). At 48h post transfection, cells were lysed in 500 μl of SDS sample buffer. In parallel, 293T cells transduced with SCRPSY-EMPTY and SCRPSY-TRIM69 were challenged with serially diluted VSV-GFP to demonstrate TRIM69 activity.

### VSIV *in vitro* evolution, RNA extraction, PCR and sequencing

MT4 cells expressing TRIM69 were seeded and induced with doxycycline (200 ng/ml) 24 h prior to challenge with FL-VSIV-GFP at low MOI (<1% infection following 16 h incubation). The level of infection based on percentage of GFP positive cells was monitored daily. Once the culture was overwhelmed, supernatant was filtered (0.45 μm) and used to infect a second culture. Following passage 2, CPE was observed 24 h postinfection (in the presence of TRIM69). The supernatant of the passaged virus was filtered (0.45 μm) and stored. Cell pellets were resuspended in TRIzol (Invitrogen) and viral RNA was isolated from infected cells using a hybrid TRIzol and RNeasy extraction (Qiagen) protocol. Viral cDNA was reverse transcribed (SuperScript III) using random hexamer primers. All VSIV coding regions were PCR-amplified and the PCR products were directly sequenced using Sanger sequencing (Eurofins Genomics).

### TRIM69 sequences collection and alignment

TRIM69 sequences were obtained from publicly available databases such as ENSEMBL and GenBank using TBLASTN (68). Supplementary table (SequenceTable.xls) lists all the accession numbers and species used. The protein sequences were aligned using MAFFT (69) and a codon alignment generated based on the protein alignment using PAL2NAL (70). The alignment was screened for recombination in HyPhy (71) using single breakpoint recombination (SBP) and Genetic Algorithm for Recombination Detection (GARD) (72).

### Phylogenetics and positive selection analyses

The best substitution model was selected using BIC in jModeltest (73) and the maximum likelihood phylogeny was reconstructed with the latter model in PhyML with 1000 bootstrap replicates (74). The primate lineage of the gene tree, which is more extensively sampled for species, was used in CODEML (38) to detect sites under positive selection.

## Acknowledgements

We thank R. Fouchier, T. Zang, P. Bieniasz, G. Towers and A. Fletcher for reagents. This study was funded by Medical Research Council (MRC) grants MR/K024752/1(to SJW), MC_UU_12014/10 (to SJW and MP), MR/P022642/1 (to SJW and SJR) and MC_UU_12014/12 (to JH) as well as Wellcome Trust support 201366/Z/16/Z (to SJR).

